## Supplementary Tables and Figures for "Enzalutamide Induces Cytotoxicity in Desmoplastic Small Round Cell Tumor Independent of the Androgen Receptor"

**Supplementary Table 1:** Antibodies for Western Blot

| Antibody | Company | Catalog # | WB Dilution |
|---|---|---|---|
| β-Actin (8H10D10) | Cell Signaling | 3700 | 1:1000 |
| AR | Cell Signaling | D6F11 | 1:1000 |
| LCK | Cell Signaling | L22B1 | 1:500 |
| EWSR1 (N-term) | Lab Created | N/A | 1:1000 |
| MERTK | Cell Signaling | 348E6 | 1:1000 |
| FLAG | Sigma | F3165 | 1:1000 |

**Supplementary Table 2:** RT-qPCR Primers

| Gene | Fwd Primer (5' -> 3') | Rev Primer (5' -> 3') |
|---|---|---|
| ACTB | GCAAAGACCTGTACGCCAAC | AGTACTTGCGCTCAGGAGGA |
| MERTK | GGAAATAGCTACGCGGGGAA | GAAAAGGTGGGGCGGTCTAA |
| FGFR4 | AGTCAAGCTCATCCCTGGTA | CAGAGTGGGTCGAGAGGTAGAT |
| EPHB3 | GCCCTTGGTGCTGTCATAA | CACACTCACACATCCCATCTC |
| CCL25 | CTTGACCCAGTGGATATCGGT | GAGCACAGCCCACCCAAT |
| AR | GGCCGAATGCAAAGGTTCTC | GCCTTCTAGCCCTTTGGTGT |
| WT1 (c-term)/ EWSR1-WT1 | CCATACCAGTGTGACTTCAAGG | TGTGGGTCTTCAGGTGGTC |

**Supplementary Table 3:** shRNA Sequences

| Target | Name | Sequence (5' -> 3') |
|---|---|---|
| AR | shAR #1 | GAGCGTGGACTTTCCGGAAAT |
| AR | shAR #2 | GATGTCTTCTGCCTGTTATAA |
| AR | shAR #3 | ACCGAGGAGCTTTCCAGAATC |
| AR | shAR #4 | AGCTGCTCCGCTGACCTTAAA |
| WT1 | shWT1 | GCAGCTAACAATGTCTGGTTA |

**Supplementary Table 4:** DSRCT Xenograft Seeding

| **First Author** | **Year** | **Cell Line** | **Cell Number** | **Mouse Type** | **Mouse Sex** | **Injection Site** | **Matrigel** |
|---|---|---|---|---|---|---|---|
| Nishio | 2002 | JN-DSRCT-1 | 5.00E+07 | SCID | Female | Subcutaneous | No |
| Hayes-Jordan | 2018 | JN-DSRCT-1 | 2.00E+06 | NOD/SCID gamma | Male | Intraperitoneal | No |
| Uboldi | 2017 | JN-DSRCT-1 | 5.00E+07 | NOD/SCID gamma | Not specified | Subcutaneous | No |
| Erp | 2020 | JN-DSRCT-1 | 5.00E+06 | SCID | Male | Subcutaneous | Yes |
| Smith | 2020 | JN-DSRCT-1 | 1.00E+07 | NOD/SCID gamma | Female | Subcutaneous | Yes |
| Smith | 2020 | BER-DSRCT | 1.00E+07 | NOD/SCID gamma | Female | Subcutaneous | Yes |
| Smith | 2020 | BOD-DSRCT | 1.00E+07 | NOD/SCID gamma | Female | Subcutaneous | Yes |
| Smith | 2020 | SK-DSRCT2 | 1.00E+07 | NOD/SCID gamma | Female | Subcutaneous | Yes |
| Smith | 2020 | BER-DSRCT | 1.00E+07 | NOD/SCID gamma | Female | Intraperitoneal | Yes |
| Smith | 2020 | SK-DSRCT1 | 1.00E+07 | NOD/SCID gamma | Female | Intraperitoneal | Yes |
| Ogura | 2021 | JN-DSRCT-1 | 1.00E+07 | NOD/SCID gamma | Female | Subcutaneous | Yes |
| Erp | 2022 | JN-DSRCT-1 | 5.00E+06 | SCID | Male | Subcutaneous | Yes |

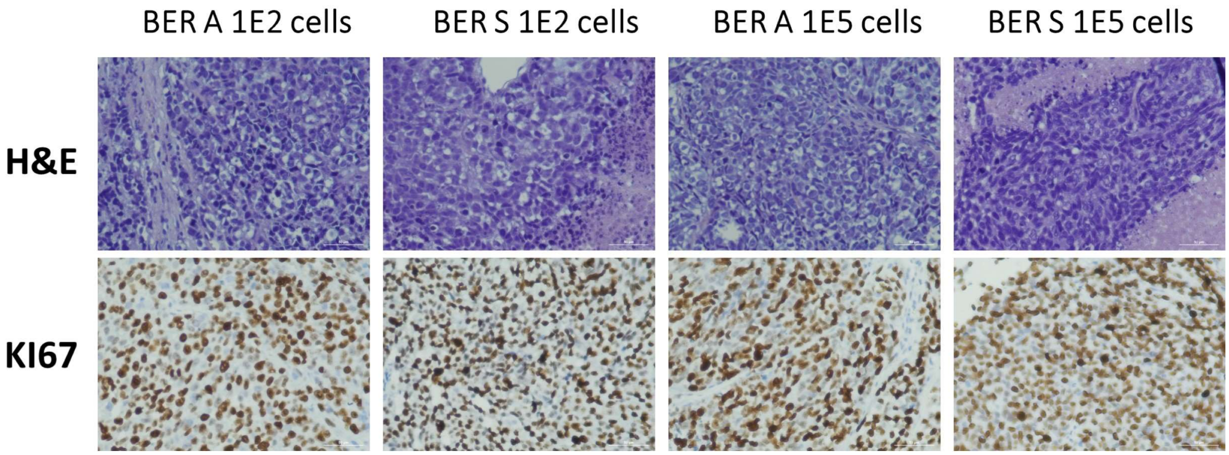


**Supplementary Fig 1. DSRCT xenograft Morphology. (A)** Representative H&E and KI67 staining of BER-DSRCT xenografts seeded from (A) adherent or (S) sphere culture cells (scale bar = 50 μm).

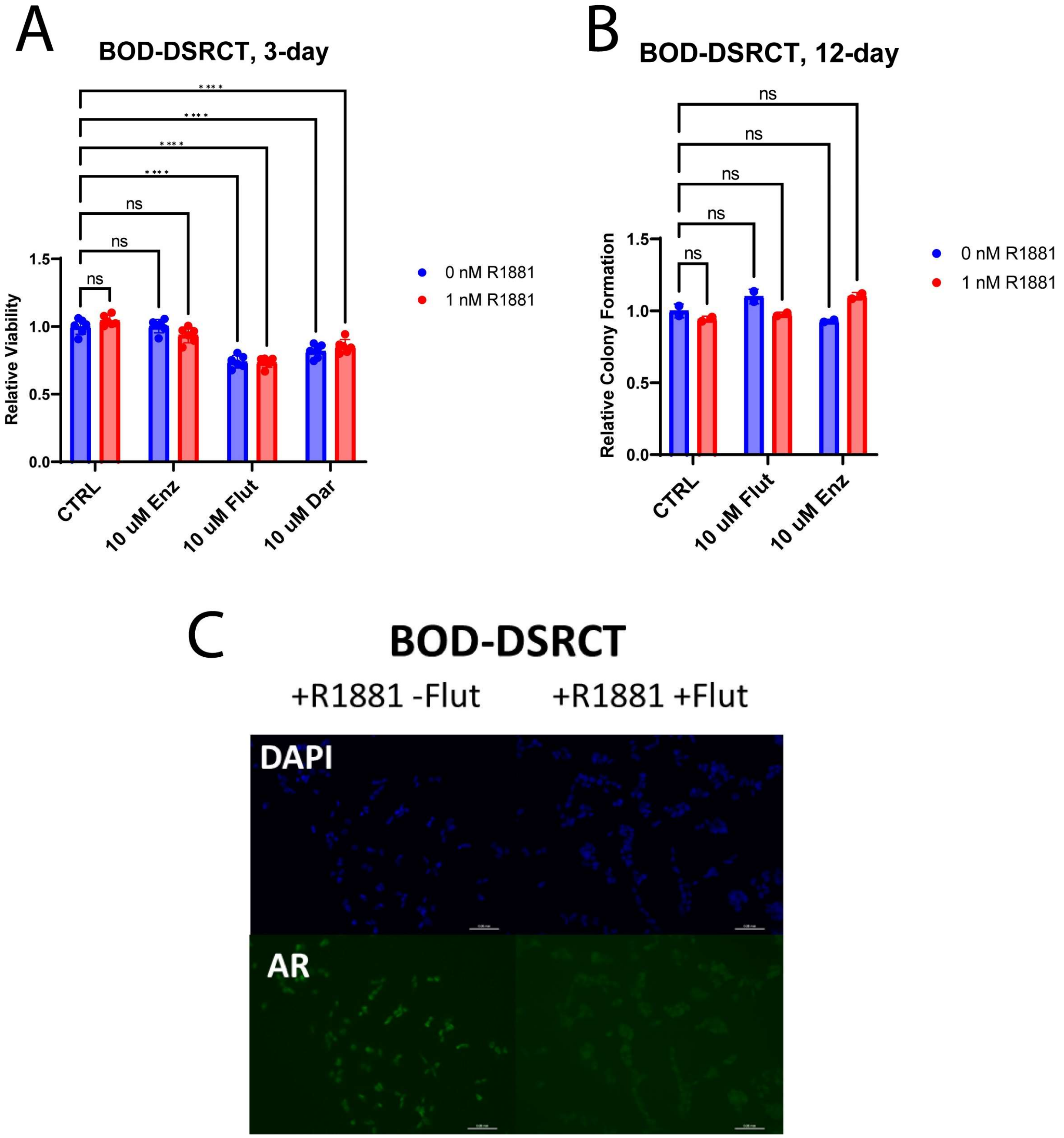


**Supplementary Fig 2. BOD-DSRCT AR Response**. **(A)** Relative viability of BOD-DSRCT cells treated with 10µM enzalutamide, flutamide, or darolutamide for 72 hrs with or without 1nM R1881 (n=3, * $p<0.05$, ** $p<0.01$, *** $p<0.001$, **** $p<0.0001$). **(B)** Colony formation assays examining the effect of flutamide and enzalutamide on DSRCT growth over a 14-day period (n=2, ns = nonsignificant). **(C)** Immunofluorescence imaging of DAPI and AR in DSRCT cells treated with 1nM R1881 and with (+) or without (-) 10µM flutamide for 24 hrs (n=2, scale bar = 50 µm).

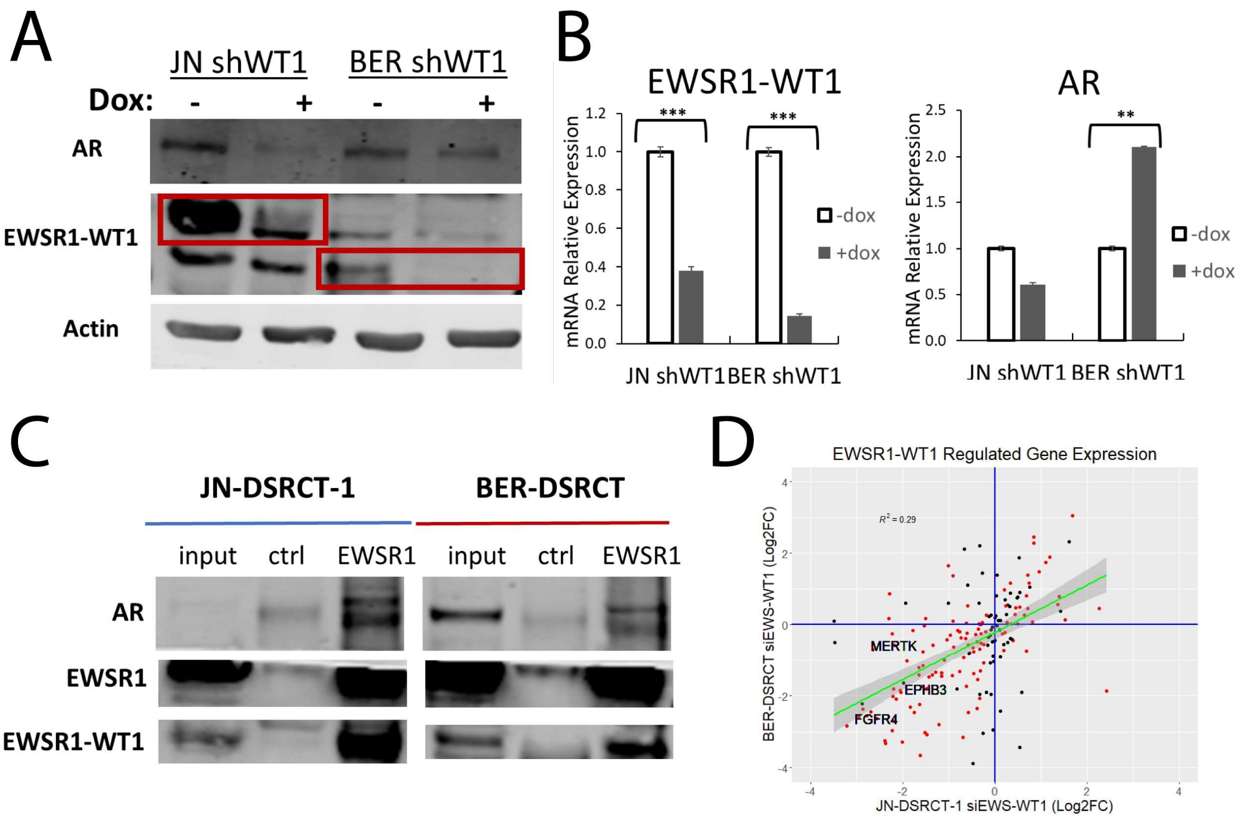


**Supplementary Fig 3. AR regulation by EWSR1-WT1**. **(A)** Western blot of EWSR1-WT1, AR, and ACTIN protein expression in shWT1 cell lines that deplete EWSR1-WT1 with dox (+) addition (representative blot, n=2). **(B)** RT-qPCR of EWSR1-WT1 and AR expression in JN-DSRCT-1 and BER-DSRCT shWT1 cell lines with or without dox (n=3, * $p<0.05$, ** $p<0.01$). **(C)** Immunoprecipitation-Western blot using anti-EWSR1 or control antibody in JN-DSRCT-1 and BER-DSRCT cells demonstrating an interaction between AR and EWSR1-WT1 and/or native EWSR1. **(D)** Scatterplot of log2FC gene expression change of AR-EWSR1-WT1 co-occupied genes with EWSR1-WT1 depletion as measured with RNA-seq.

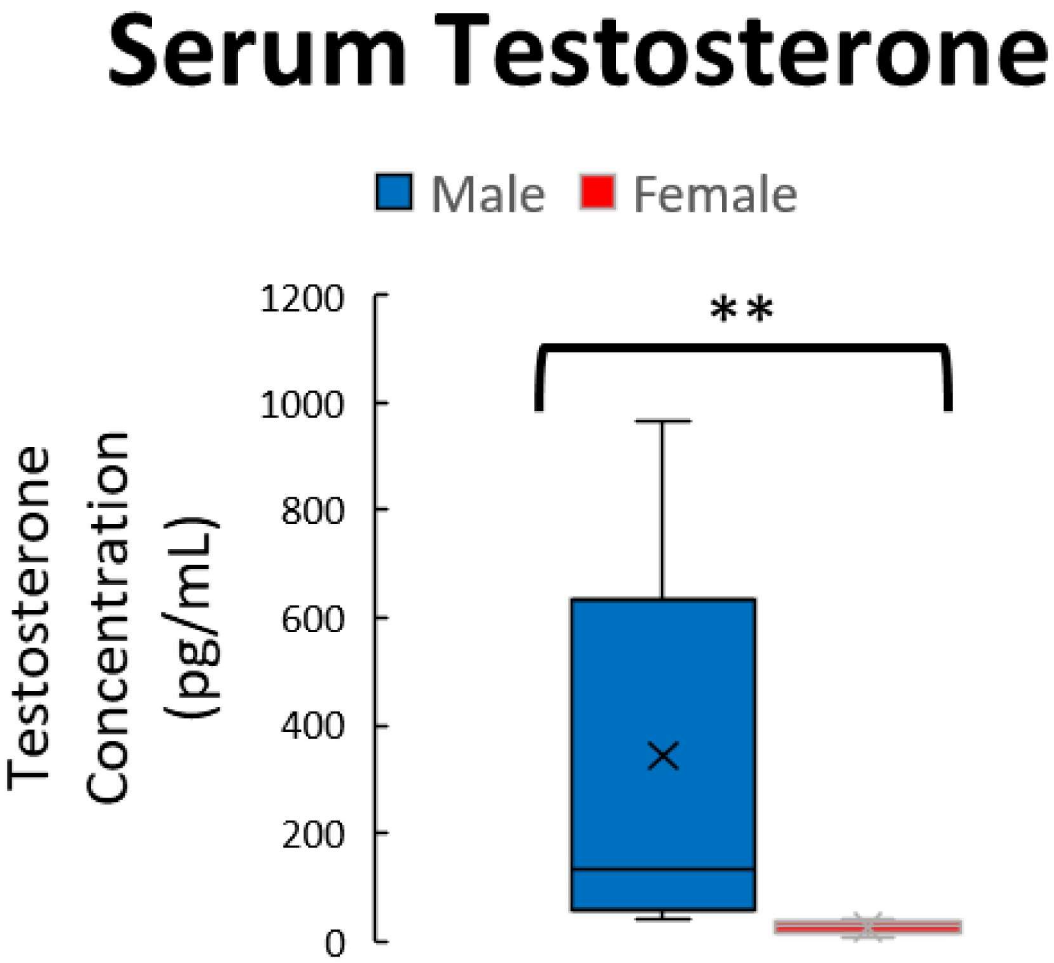


**Supplementary Fig 4. Mouse testosterone differences with sex**. Testosterone concentration in the blood of male and female mice as measured by ELISA (n=8, ** p<0.01).

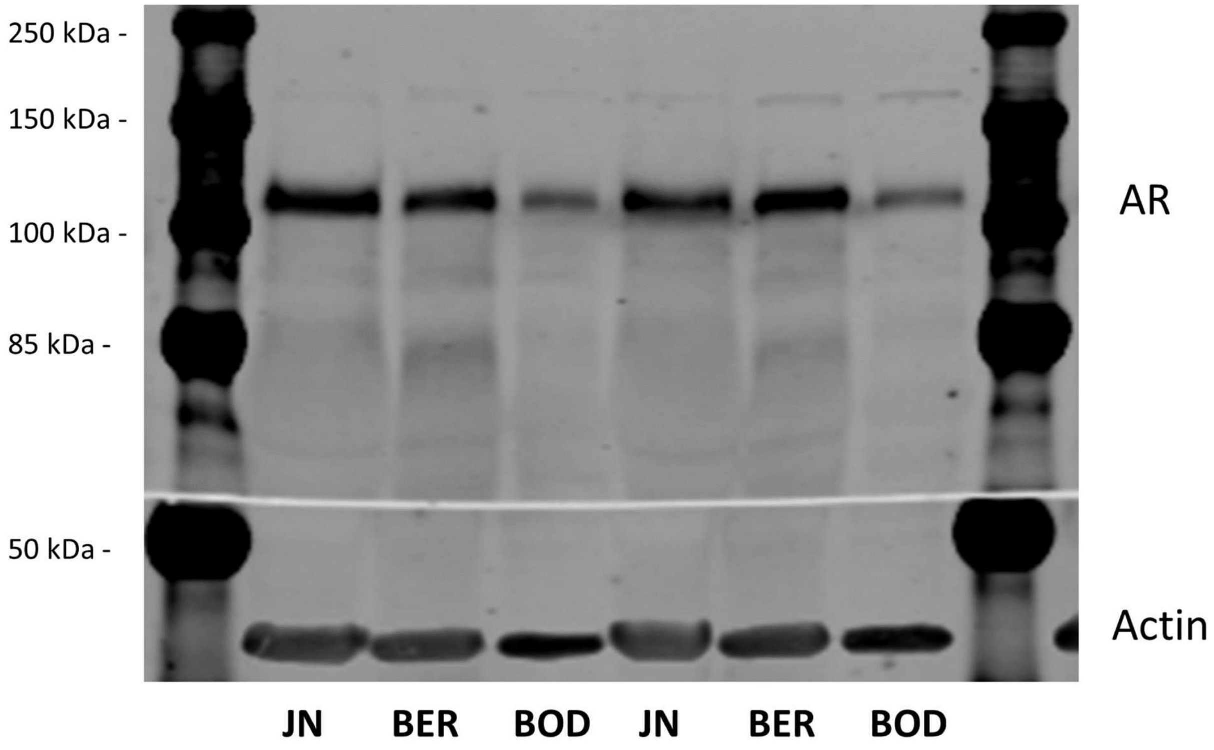


**Supplementary Fig 5. Lack of AR variants in DSRCT.** Western blot of AR in three DSRCT cell lines examining 50 to 250kDa region for native AR and/or variants. Only native AR was detected.

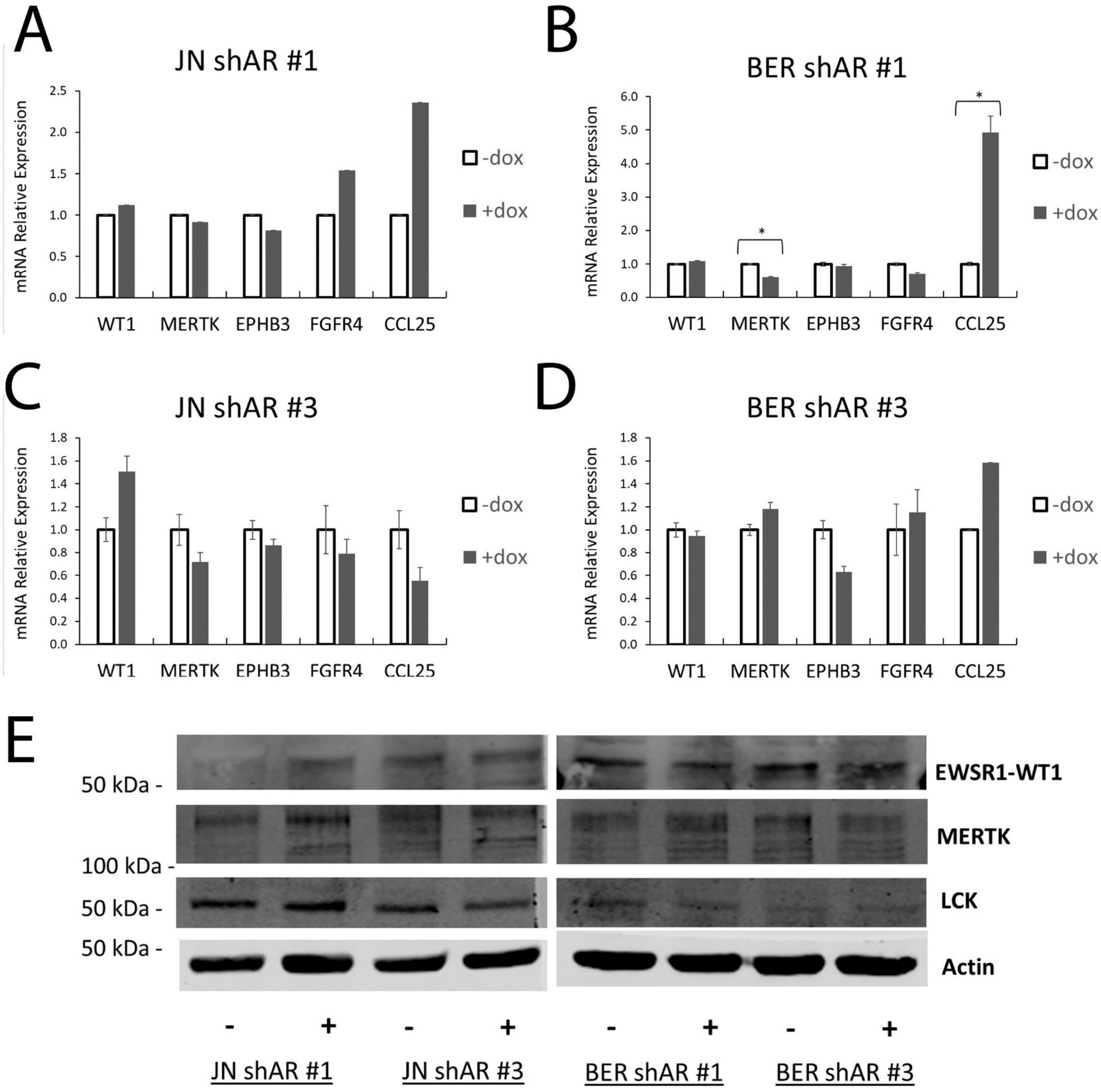


**Supplementary Fig 6. AR depletion Gene Expression. (A-D)** RT-qPCR of *EWSR1-WT1* (measured with WT1 C-term primers), *MERTK, EPHB3, FGFR4*, and *CCL25* gene expression in **(A)** JN-DSRCT-1 shAR #1, **(B)** BER-DSRCT shAR #1, **(C)** JN-DSRCT-1 shAR #2, and **(D)** BER-DSRCT shAR #2 cells lines with (+) or without (-) dox addition to deplete AR. **(E)** Western blot examining EWSR1-WT1, MERTK, LCK, and ACTIN in JN-DSRCT-1 and BER-DSRCT shAR #1 and 3 cell lines treated for four days with (+) or without (-) dox to deplete AR (n=3).

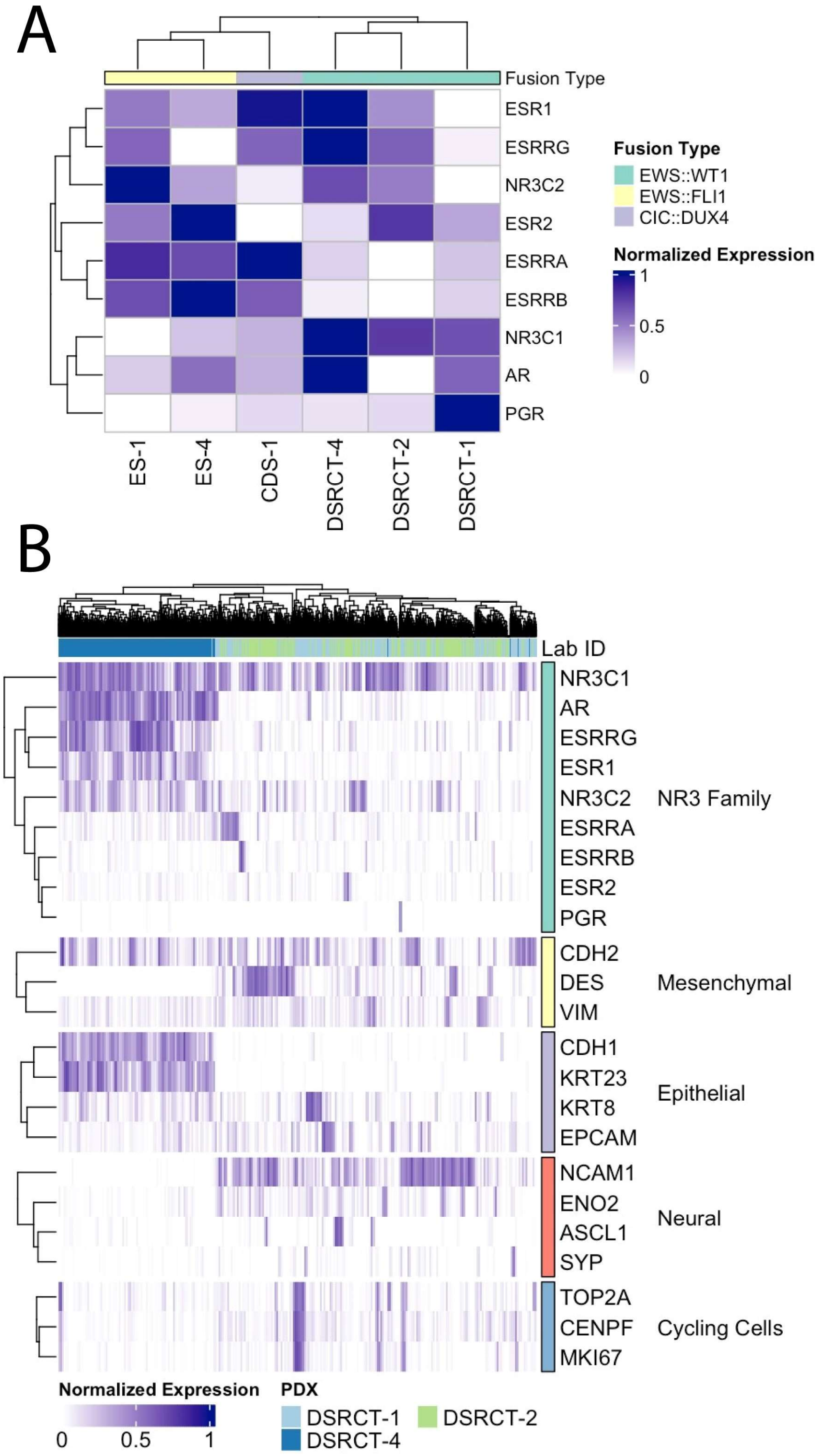


**Supplementary Fig 7. NR3 nuclear receptor expression in DSRCT. (A)** Heatmap of NR3 nuclear receptor expression from bulk RNA-seq of patient-derived xenografts from DSRCT (n=3), Ewing sarcoma (ES, n=2), and CIC-DUX4 (CDS, n=1). **(B)** Heatmap from snRNA-seq of three DSRCT patient-derived xenografts showing expression of NR3 nuclear receptors, mesenchymal markers, epithelial markers, neuronal markers, and markers of cell cycling.
